# Rapid Audiovisual Integration Guides Predictive Actions

**DOI:** 10.1101/2022.12.09.519832

**Authors:** Philipp Kreyenmeier, Anna Schroeger, Rouwen Cañal-Bruland, Markus Raab, Miriam Spering

## Abstract

Natural movements, such as catching a ball or capturing prey, typically involve multiple senses. Yet, laboratory studies on human movements commonly focus solely on vision and ignore sound. Here we ask how visual and auditory signals are integrated to guide interceptive movements. Human observers tracked the brief launch of a simulated baseball, randomly paired with batting sounds of varying intensities, and made a quick pointing movement at the ball. Movement endpoints revealed systematic overestimation of target speed when ball launch was paired with a loud versus a quiet sound, even though sound was never informative. This effect was modulated by the availability of visual information: sounds biased interception when ball presentation time was short. Continuous eye movements uncovered that sound affected interception within 125 ms of ball launch and revealed a correction of the sound-induced bias when more visual information was available. Our findings suggest that auditory and visual signals are integrated to guide interception and that this integration process must occur early at a site that receives auditory and visual signals within an ultrashort timespan.

## Significance Statement

Almost all everyday actions, from catching a ball to driving a car, rely heavily on vision. Even though moving objects in our natural visual environment also make sounds, the influence of auditory signals on motor control is commonly ignored. This study investigates the effect of sound on vision-guided interception. We show that sound systematically biases interception movements, indicating that observers associate louder sounds with faster target speeds. Using eye movements as a continuous indicator of multisensory processing, we demonstrate that vision and sound are rapidly integrated within 125 ms of an object’s appearance. Training and rehabilitation approaches in sports and medicine could harness the finding that interceptive hand movements are driven by multisensory signals and not just vision alone.

## Rapid Audiovisual Integration Guides Predictive Actions

When intercepting a rapidly moving object with our hands—swatting a fly or catching a ball—we rely heavily on vision. Humans and other animals direct their eyes at moving objects of interest to sample critical visual information and to increase performance accuracy [1–5]. However, other sensory modalities also supply information that might be used to guide behaviour in interception tasks. Indeed, in goalball—an interceptive sport for visually-impaired athletes—players rely solely on auditory information to locate and intercept a ball [6]. Our study addresses the question whether and under which conditions vision-guided interceptive actions rely on sound information in normally-sighted observers.

In our natural environment, object motion is almost always accompanied by sound, which can alter visual motion judgements [7–11]. For instance, hitting a ball with a bat or racket creates an impact sound, and its volume provides information about hit intensity and launch speed. Accordingly, impact sounds can bias perceived ball-bounce locations and perceptual ball speed judgements, suggesting that observers use auditory information to predict ball trajectories [12,13]. When integrating information from different modalities, Bayesian models of multisensory integration predict that sensory signals are combined by weighing them according to their uncertainty [14–17]. Therefore, auditory signals may bias visual perception most in tasks with visual uncertainty resulting from ambiguity (e.g., visual illusions [18]), or sparseness (e.g., poor viewing conditions).

Our study probes this interaction between visual certainty and auditory cues in a real-world inspired, fast-paced movement interception task **(Fig.1A)** during which observers track the brief launch of a simulated baseball moving across the screen and intercept it at a predicted location with a quick pointing movement. We manipulated sound intensity of the simulated ball launch and visual certainty by varying the presentation time of the ball. In this challenging task, we measured observers’ eye and hand movements towards the ball as indicators of their abilities to estimate its speed and to predict its trajectory. This approach allows us to directly test whether principles of Bayesian audio-visual integration, previously demonstrated for pure perceptual tasks, apply to predictive action control. We hypothesized, first, that auditory cues would systematically bias ball speed estimation. Specifically, we expected that observers would overestimate speed when ball launch was accompanied by a loud batting sound (indicating a harder hit and higher launch speed) as compared to a quiet batting sound (indicating a softer hit and lower launch speed). Second, we expected that the auditory cue’s influence would scale with visual certainty, implying that observers rely more on the auditory cue when visual presentation times are short, in line with the predictions of a Bayesian approach to multisensory cue combination (**Fig.1B**). Further, measuring continuous eye movements during this track-intercept task allows us to investigate the temporal dynamics of audio-visual integration and the extent to which observers rely on auditory information over time.

**Figure 1.**
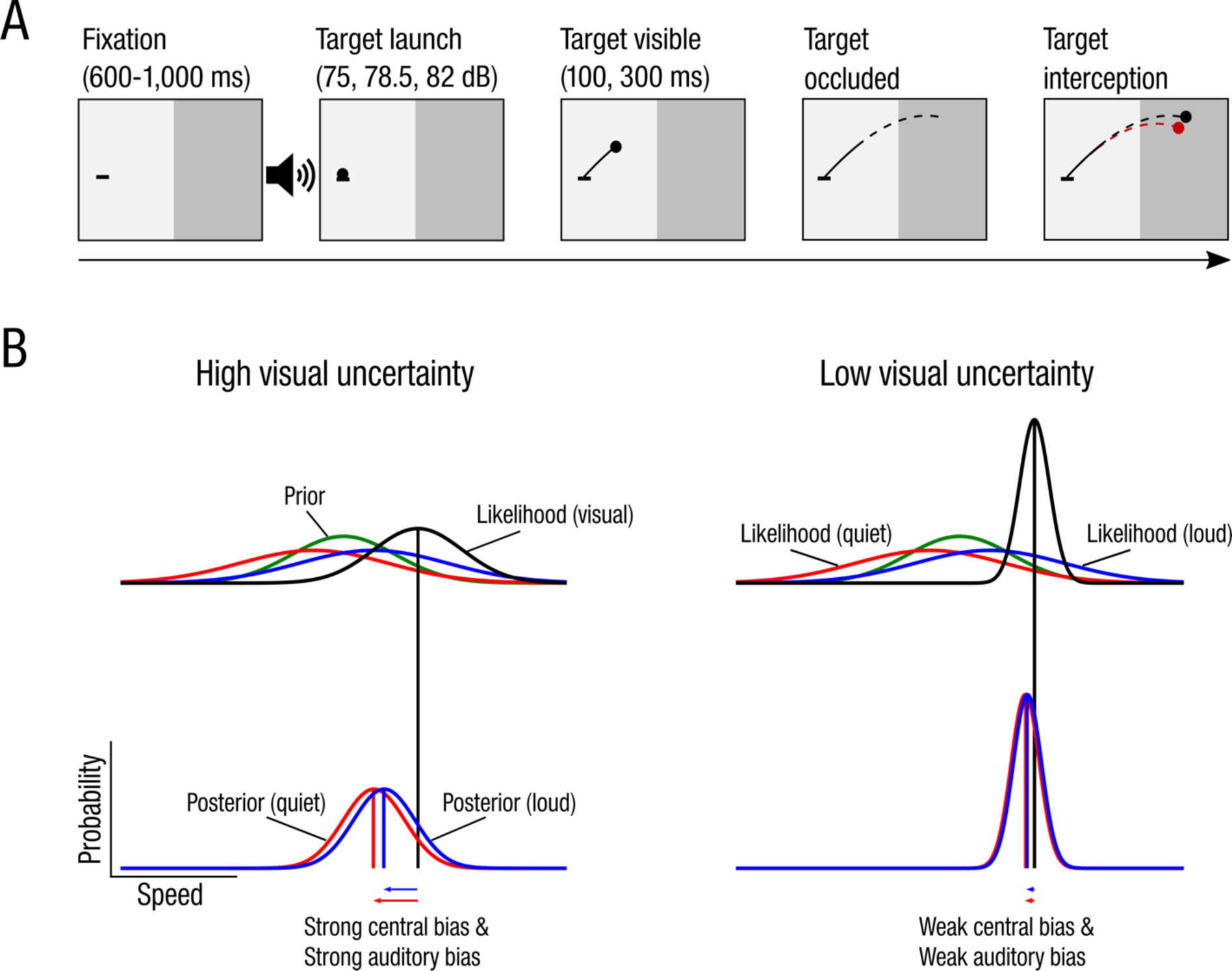
**(A)** Timeline of a single trial. After initial fixation, observers tracked the launch of a simulated batted baseball (black dot) moving along one of five trajectories, defined by different launch speeds. Ball launch was randomly paired with batting sounds of varying intensities. After a short presentation time (solid black line) the ball disappeared (dashed black line), and observers had to predict its trajectory and intercept it within a ‘hit zone’ (dark grey area). Feedback of interception position (red disc) and actual target position (black disc) at the time of interception was presented at the end of each trial. Far right panel illustrates the analysis of estimated target speed: The black lines show the actual target trajectory in a given trial. The hypothetical target trajectory (dashed red line) that produced the smallest Euclidian distance to the interception position (red disc) is labelled estimated target speed for this trial. **(B)** Predictions of a Bayesian model with a central prior (green curve) around the mean target speed. Auditory cues had no diagnostic information and are assumed to be highly uncertain (wide auditory likelihood functions; red and blue curves). Uncertainty of visual estimates (black curves; illustrated for a high target speed) is reflected in the width of the likelihood functions. Under high visual uncertainty (left), posteriors (bottom left) will be strongly biased by the prior and auditory cues. Under low visual certainty (right), posteriors will only be marginally shifted toward the mean and show almost no influence of auditory cues.

## Results

Observers (*n*=16) tracked the brief launch of a simulated baseball moving at different speeds and randomly paired with batting sounds of varying intensities **(Fig.1A)**. The ball disappeared shortly after launch, either after 100 or 300 milliseconds (ms), resulting in conditions of low (short presentation condition) and high (longer presentation condition) visual certainty. Observers then intercepted the ball with a quick pointing movement along the predicted trajectory within a ‘hit zone’ (**Fig.2A**).

**Figure 2.**
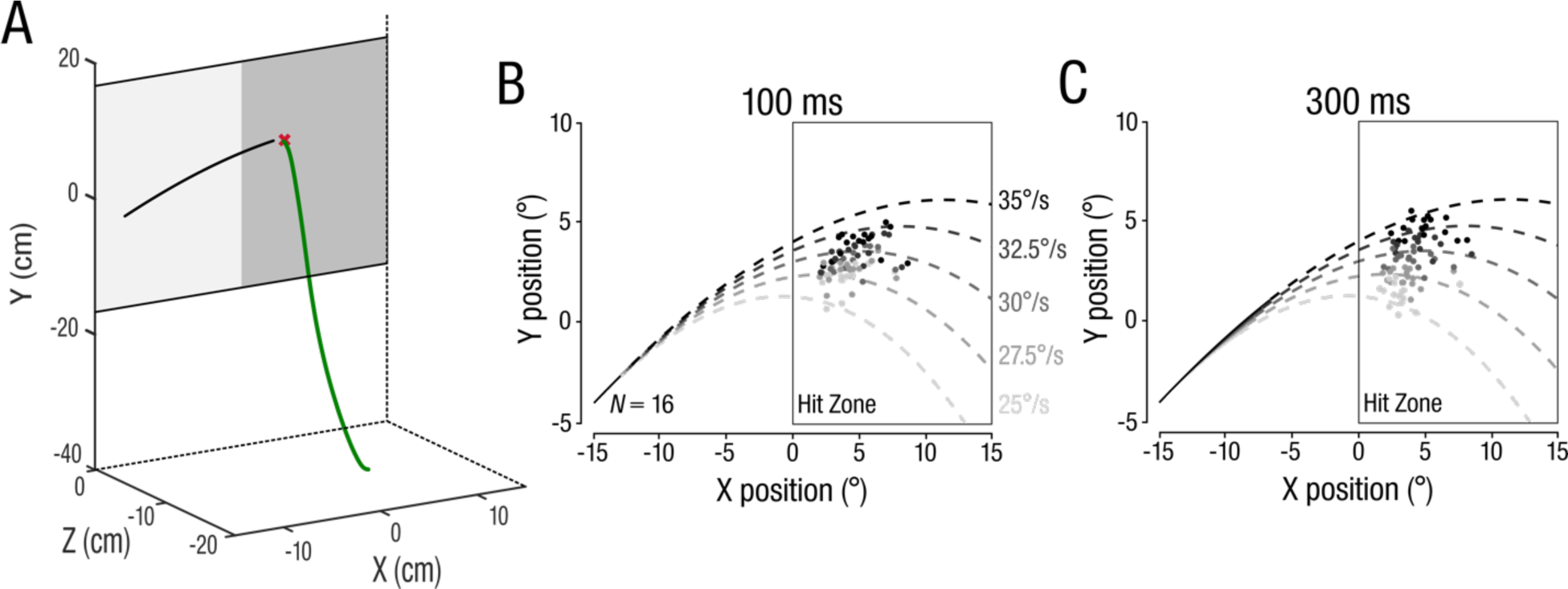
**(A)** Example of a hand (green) position trace. Black line represents the 2D target position and red cross indicates the interception position. **(B-C)** Mean individual observer 2D interception positions for the **(B)** 100 ms and **(C)** 300 ms presentation durations. Each data point indicates one observer’s mean interception position per each of the five target speeds.

In our task, observers had to rely on visual information of speed during the ball’s brief presentation to extrapolate and intercept it accurately. Mean 2D interception positions show that observers intercepted targets along their predicted trajectories and discriminated between different target trajectories in both the 100-ms (**Fig.2B**) and 300-ms conditions (**Fig.2C**). However, interception endpoints strongly regressed toward the intermediate trajectory in the 100-ms condition, indicating that observers were uncertain about the target trajectory. In contrast, in the 300-ms condition, observers intercepted balls more accurately along their trajectories (**Fig.2C**).

## Auditory Cues Bias Target Speed Estimates when Visual Information is Uncertain

Bayesian observer models of multisensory cue combination predict that prior knowledge and sensory cues are weighed according to their uncertainty (**Fig. 1B**). Assuming a central prior that biases visual evidence toward the mean target speed and that auditory cues induce systematic biases, we can derive two critical predictions from these models: Under high visual uncertainty, target speed estimates will be strongly influenced by the prior (regression toward the mean) and auditory cues (systematic bias). Conversely, under low visual uncertainty, resulting speed estimates will only be marginally shifted toward the prior and the influence of the auditory cue becomes negligible. To test these predictions, we used observers’ target speed estimates as the primary outcome measure. We determined the estimated target speed from the 2D interception position by finding the hypothetical trajectory that produces the smallest Euclidian distance to the interception position (dashed red line in **Fig.1A**). **Figure 3** shows observers’ estimated target speed as a function of physical target speed, separately for each sound volume. If speed estimates were accurate, they would fall along the diagonal (dashed line). To test the prediction of a stronger prior influence on target speed estimates under high versus low visual uncertainty (**Fig.1B**), we ran a linear mixed model with *physical target speed* and *presentation time* as predictors.

**Figure 3.**
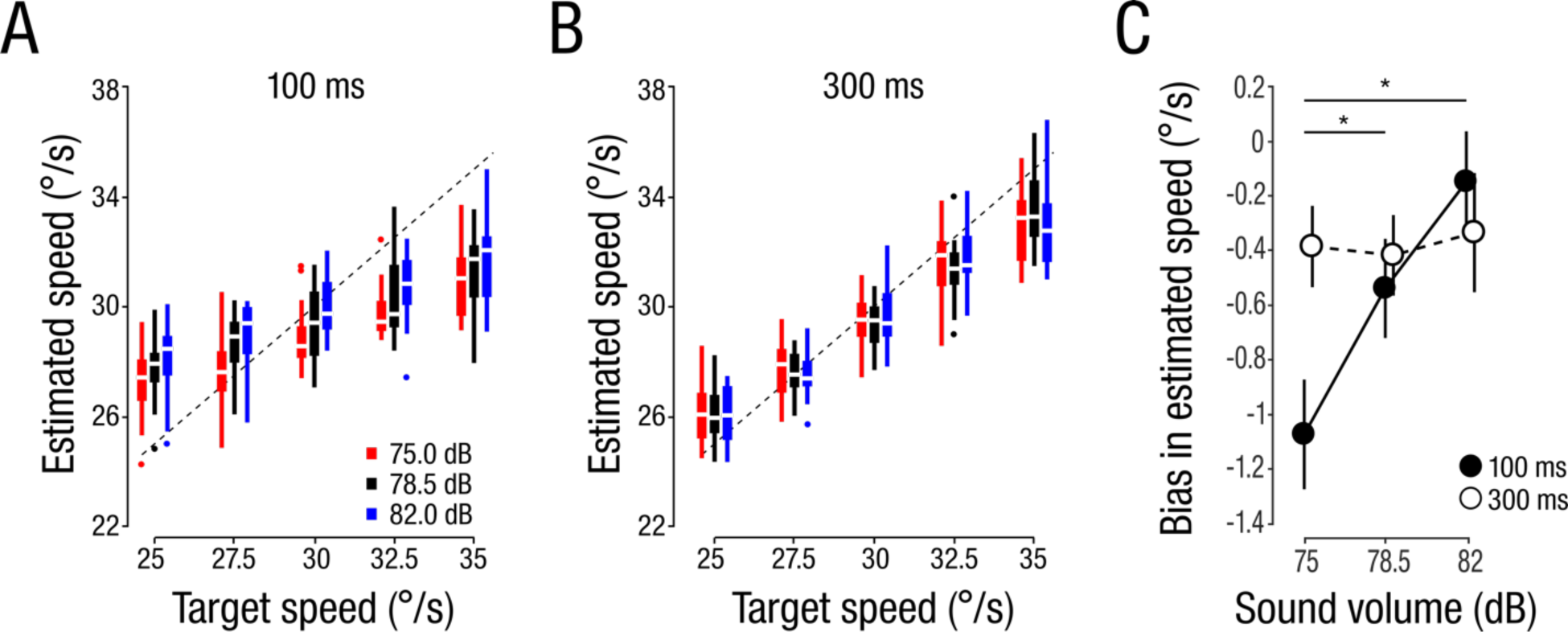
(**A-B**) Boxplots of estimated target speed (*n*=16) as a function of physical target speed. Colours denote sound volume conditions and dashed lines indicates veridical estimates. (**B**) 100 ms condition. (**C**) 300 ms condition. (**C**) Effect of stimulus volume on the bias in the estimated target speed averaged across target speeds, separately for the 100 ms (filled circles) and 300 ms (open circles) condition. Circles and error bars denote the mean ±1 within-subject standard error of the mean (SEM). Asterisks denote significant post-hoc comparisons (*p*<.05).

Physical target speed was a significant predictor of estimated target speed for both presentation times (100 ms: *β*=.37; *t*(15)=9.66; *p*<.001; 300 ms: *β*=.72; *t*(15)=14.94; *p*<.001). In line with our predictions, we found a significant difference between slopes for the 100 and 300-ms conditions (*β*=-.35; *t*(15)= −15.92; *p*<.001), confirming that observers’ speed estimates regressed more toward the mean (indicating a stronger influence of the prior) in the 100-ms condition, as compared to the 300-ms condition.

Next, we assessed whether and under which conditions sound volume influenced visual target speed estimates. We hypothesized that sound volume would systematically bias observers’ target speed estimates and that this bias would depend on the certainty of the visual speed signal. Accordingly, we found that observers systematically underestimated target speed when ball launch was paired with a quiet batting sound and overestimated target speed when the ball was paired with a loud batting sound. This effect was consistent across all target speeds at short presentation duration (**Fig.3A**). Conversely, at long presentation duration, sound volume did not systematically affect estimated target speed (**Fig.2B**). To assess the differential effects of sound volume depending on presentation duration, we calculated each observer’s bias in speed estimation across target speeds (mean difference between estimated and physical target speed; **Fig.3C**). A 2 × 3 rmANOVA with factors presentation time and sound volume revealed a significant main effect of sound volume (*F*(2,30)=4.91, *p*=.029, *η_p_^²^*=0.25) and no main effect of presentation time (*F*(1,15)=0.60, *p*=.45, *η_p_^²^*=0.04). A significant interaction between presentation time and sound volume (*F*(1.43,21.47)=20.28, *p*<.001, *η_p_^²^*=0.57) confirms the profound effect of auditory cues on manual interception when visual information is sparse, but not when the target is presented sufficiently long to base speed estimation for interception on visual information alone. These findings show that when visual information was sparse and thus uncertain, target speed estimates were strongly biased toward the mean and systematically influenced by the auditory cue. Conversely, when visual uncertainty was low, estimated target speeds almost perfectly scaled with physical target speed (weak influence of the prior) and showed no impact of auditory cues. These findings are aligned with predictions of a Bayesian model of multisensory cue combination that weighs prior information and sensory cues according to their uncertainty.

## Eye Movements Reveal Temporal Dynamics of Audio-Visual Integration

The extent to which observers relied on sound depended on the certainty of the visual speed signal—i.e., presentation duration (low certainty for short, high certainty for longer presentations). The impact of the auditory signal decreased with increasing presentation duration. To assess how differences in auditory cue use in the long and short presentation time conditions unfolded over time, we analysed observers’ continuous eye movements during the interception task.

Observers tracked the simulated baseball with their eyes using a combination of smooth pursuit and saccadic eye movements (**Fig.4A**). They typically made an early catch-up saccade shortly after target onset (124.69 ±38.18 ms; mean ± SD). Subsequent predictive saccades were made after target disappearance to the predicted interception location. Eye movement endpoints, based on the 2D eye position at the time of interception, reflect observers’ ability to estimate target speed. **Figure 4B** shows that observers underestimate target speed in the presence of a quiet sound and overestimate speed when paired with a louder sound, akin to observations for manual interception responses (**Fig.2C**). Accordingly, target speed estimates based on eye and hand movement endpoints were strongly correlated on a trial-by-trial basis with a median correlation of *r*=0.70 (**Fig.4C**).

**Figure 4.**
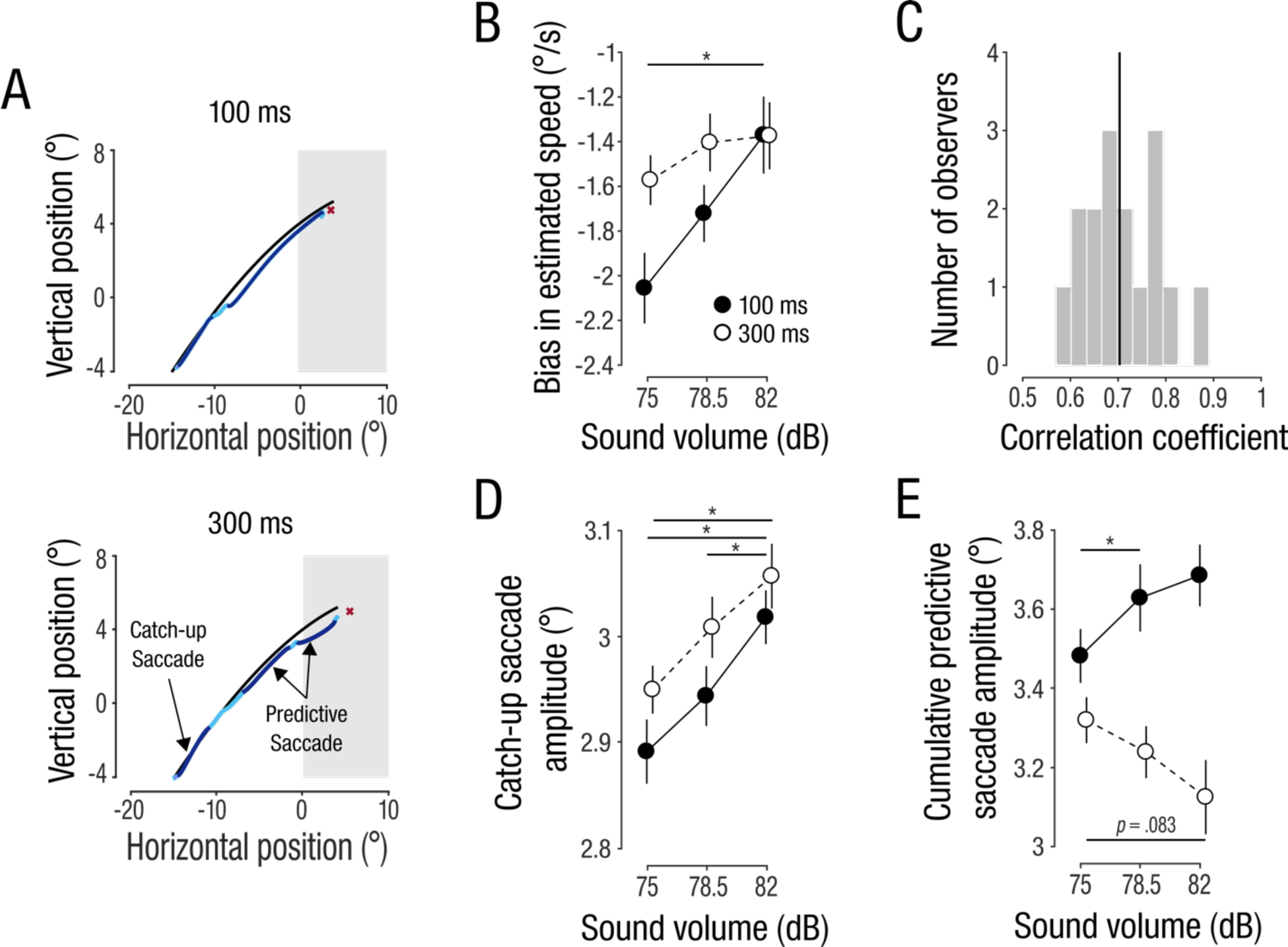
(**A**) 2D eye position traces of two representative trials. Bright blue segments are smooth pursuit, eyes’ continuous tracking of moving targets, dark blue segments are saccades. (**B**) Effect of sound volume on estimated target speed based on final eye position. (**C**) Histogram of trial-by-trial correlation coefficients from all observers. Black line indicates median across observers. (**D**) Effect of sound volume on vertical saccade amplitudes for first catch-up saccades after target onset. (**E**) Cumulative vertical amplitudes of all subsequent predictive saccades. Circles indicate means across observers; error bars are ±1 within-subject SEM. Asterisk denotes significant post-hoc comparison (*p*<.05).

We next assessed whether eye movements reveal the time course of audio-visual integration by analysing the vertical amplitude of the first catch-up saccade (early integration; initiated ∼125 ms after target onset) and the combined vertical amplitude of subsequent, predictive saccades (later integration). For this analysis, we excluded trials where the first catch-up saccade was made in anticipation of target onset (<50 ms latency; 3.92% of trials). We found an influence of sound volume at an early timepoint (main effect of sound volume: *F*(2,30)=29.04; *p*<.001; *η_p_^²^*=0.66; **Fig.4D**), consistently observed across physical target speeds (**Fig.S1**). Neither the main effect of presentation time nor the interaction term was significant (all *p*>.487), indicating that the influence of the auditory cue occurred before any differences in presentation duration could impact visual speed judgments. Note that the first catch-up saccade not only showed consistent and similar effects of sound volume between presentation times but also scaled similarly with physical target speeds (**Fig.S1**). This further indicates that early catch-up saccades were finely tuned to the sensory properties of the target and were programmed before differences between presentation durations emerged.

By contrast, predictive saccades that occurred later in the trial had increased amplitudes with increasing sound volume in the 100 ms condition but had decreased amplitudes for the longer presentation duration (**Fig.4E**). This finding was consistent across target speeds (**Fig.S1**) and confirmed by a significant sound volume × presentation time interaction effect (*F*(2,30)=20.85; *p*<.001; *η_p_^²^*=0.58). A significant main effect of presentation time (*F*(1,15)=9.78; *p*=.014; *η_p_^²^*=0.39) is likely due to smaller saccade sizes in the 300-ms condition, which generally elicits stronger pursuit; the main effect of sound volume was not significant (*F*(1.38,20.75)=0.60; *p*=.501; *η_p_^²^*=0.04).

These opposing effects in the 100- and 300-ms condition indicate a correction of the early auditory influence with the availability of additional visual information. This was further supported by the finding that predictive saccades in the 300 ms condition scaled more with physical target speeds than predictive saccades in the 100 ms condition (**Fig.S1**).

## Discussion

Predicting the trajectory of a moving object is a fundamental ability that allows us to accurately hit, catch, or otherwise intercept targets [19]. Most research on interception focuses solely on vision to form trajectory predictions and guide interceptive hand movements [5,20]. Yet, in our natural environment, object motion is typically accompanied by sounds that can provide additional information about an object’s motion. Here we show that auditory signals are used in combination with visual motion information to estimate target speed for interceptive actions. Using a rapid track-intercept task in which a visual trajectory was paired with batting sounds of varying intensities we present three key findings: (1) Sound intensity of bat-ball contact systematically influences interception responses, extending well-known effects of audio-visual integration on perception to interceptive actions. (2) Congruent with Bayesian models of multisensory cue combination, integration of auditory cues and visual information depends on the certainty of the visual signal: Auditory cues influence target speed estimates only when visual information is sparse. (3) Audio-visual integration occurred within the first 125 ms of stimulus presentation and is corrected later with the availability of additional visual information. The temporal dynamics of audio-visual integration was revealed by analyzing continuous eye movements during this task. In our experiment, sound intensity was never informative of physical target speed, precluding the possibility that our results were solely caused by learning to associate certain sound volumes to certain target trajectories. Instead, our findings likely reflect a natural association between sound volume and relative target speed gained through lifelong experience. Under similar environmental conditions, louder sounds will typically correspond to higher target speeds. When splitting our data between first and second halves of the experiment, we found that the auditory influence was stronger during the first half of the experiment (**Fig.S2**). This indicates a strong association between sound volume and target speed that decreased with more experience in the task where sound volume and target speed were not associated. Together, these findings highlight the important contribution of auditory cues for vision-guided actions, particularly in situations where visual information is sparse or highly uncertain (e.g., in poor viewing conditions). These results build on a long line of literature on audio-visual signal integration for perceptual tasks [21]. The novelty of our findings lies in discovering how auditory information contributes to vision-guided interception, a fundamental ability for everyday interactions.

By manipulating the visual presentation duration of the target, we revealed that the use of auditory cues critically depends on the uncertainty of the visual motion signal. This finding fits with predictions of a Bayesian model of multisensory cue combination that weighs prior information and sensory evidence according to their uncertainty [14–17] (illustrated in **Fig. 1B**). Such a model makes two critical predictions: First, when visual information is highly uncertain, speed estimates are strongly biased by prior knowledge, in line with our findings. Our results show a strong regression toward the mean in the 100 ms condition, likely reflecting the use of a prior based on the statistics of the stimuli used in our experiment [22–24]. Interestingly, recent human neuroimaging studies related such central biases in temporal estimation tasks to activity in the hippocampal formation [25]. Alternatively, a prior might have been derived from statistics of our natural environment. Studies on visual [26] and auditory motion perception [27] revealed that observers typically rely on a slow-motion prior, in line with our finding that observers generally undershot target trajectories (see **Fig. 3**). Second, Bayesian models predict that the use of the auditory cue also depends on the uncertainty of the visual speed signal. In line with this prediction, we found that observers’ target speed estimates were only influenced by auditory cues when visual information was sparse and ignored the auditory cue when sufficient visual information was provided.

Our interception task was inspired by baseball. We used a visual target that moved along a simulated batted baseball and a naturalistic batting sound. Bayesian estimation models have recently been applied to real-world Major League Baseball data to elucidate use of prior knowledge and visual cues baseball batters rely on—e.g., a pitcher’s posture and hand position—when estimating where to swing [28]. Simple cues and heuristics are critical in baseball, where hitters only have a few hundred milliseconds to decide whether and where to swing. In this or similar rapid-decision making contexts, auditory cues may provide a critical advantage, because combining them with visual cues can reduce uncertainty [14]. Yet, future studies are needed to assess whether athletes rely on auditory cues of bat-ball contacts, in addition to prior knowledge and visual signals during real-world interceptive sports, as our findings suggest.

## Eye Movements as Sensitive Indicators of Audiovisual Integration

Eye movements are a natural, instinctive behavior in tasks that require fine-motor interactions with a visual object. When manually intercepting, hitting, or catching an object, observers track its trajectory until the point of interception [5,29]. The continuous nature of these movements provides an opportunity to relate their kinematics to ongoing cognitive task processes, such as decision-making [30]. Here we used observers’ continuous eye movements to probe the temporal dynamics of audio-visual integration. We observed a systematic influence of the auditory cue on the first catch-up saccade, initiated, on average, 125 ms after target onset. At this early time point, louder sound volumes evoked larger saccade amplitudes. When additional visual information was available (long presentation duration), subsequent saccades corrected for this early auditory effect. This finding suggests that the integration of auditory and visual signals can occur at a very short timescale, in line with findings showing early effects of audio-visual cues on pupil dilation and simple saccadic decision making [31]. Previous studies have identified the superior colliculus—a midbrain structure that is also involved in the control of eye movements [32]—as a key hub of audio-visual integration [33]. Visual and auditory signals reach this brain structure within 80 ms [34], making this area an excellent candidate for short-latency audio-visual integration. In parallel, visual and auditory signals could also be integrated in cortical sensory areas such as the middle temporal cortex, an area traditionally dedicated to early visual motion processing [35].

We conclude that auditory signals significantly and systematically impact vision-guided interceptive actions. This influence was strongest when visual information was sparse. We show that non-invasive, time-sensitive eye movement measurements can provide new behavioural evidence for early and rapid integration of auditory and visual signals.

## Materials and Methods

### Participants

We show data from 16 healthy adults (25.5 ±4.7 years; eleven females; two authors). This sample size was determined using an a priori power analysis in G*Power [36] (*power*=0.80; *alpha*=0.05) with an effect size of *η_p2_*=0.34 (main effect of stimulus volume on estimated target speed) derived from pilot data. All observers had normal or corrected-to-normal visual acuity. Study protocols were approved by the university Behavioural Research Ethics Board. Observers were compensated at a rate of $10/hour.

### Apparatus

The experimental setup combined a high-resolution stimulus display with eye and hand tracking. Display and data collection were controlled by a PC (NVIDIA GeForce GTX 1060 graphics card) using Matlab and the Psychophysics and Eyelink toolboxes [37,38]. Stimuli were back-projected onto a 41.8×33.4 cm translucent screen with a PROPixx video projector at a resolution of 1,280×1,024 pixels (120 Hz; VPixx Technologies, Saint-Bruno, QC, Canada). Two speakers (S-0264A; Logitech, Newark, CA, USA), located 40 cm to the left and right of the screen centre displayed the sound. Observers viewed stimuli binocularly at a distance of 44 cm while their right eye was recorded with an Eyelink 1000 Tower Mount eyetracker (1 kHz; SR Research, Kanata, ON, Canada). The 3D position of each observer’s right index finger was recorded with a 3D Guidance trakSTAR (120 Hz; Ascension Technology, Shelburne, VT).

### Stimuli and Procedure

In each trial, we displayed a small black disc that moved along a parabola, simulating the trajectory of a batted baseball impacted by gravity, Magnus effect, and aerodynamic drag force [for details, see ref. 39]. The ball was launched at one of five speeds, resulting in five unique trajectories (**Fig.1C-D**). The screen was separated into two zones by varying background luminance; the darker right side served as the “hit zone”, in which observers were asked to intercept the ball (**Fig.1A**). The sound of a baseball hitting a wooden bat was retrieved from a free online sound library [40] (44,100 Hz) and played at one of three volumes (75, 78.5, 82 dB) at the time of ball launch (duration of ball-bat contact sound was approx. 50 ms).

Each trial began with a random-duration fixation on a line segment that marked the ball-launch position (**Fig.1A**). The ball then moved for either 100 or 300 ms before disappearing from view. Observers were instructed to manually intercept the ball anywhere along its occluded trajectory within the hit zone. Upon interception, a red dot, indicating the finger’s interception location, and a black dot, showing actual ball position at interception, provided feedback for the observer.

Observers performed nine practice trials to familiarize themselves with the task. Batting sounds, presentation times, and target speeds were pseudo-randomly selected for each trial. The experiment consisted of 420 trials total, divided into seven blocks of 60 trials each (14 trials per condition).

### Eye and Hand Movement Recordings and Analyses

Eye and hand movement data were pre-processed offline. Filtered eye movement traces (second-order Butterworth filters with 15 Hz [position] and 30 Hz [velocity] cut-off frequencies) were aligned offline to the target start position. Saccades were detected when five consecutive frames exceeded a fixed velocity criterion of 30°/s; saccade on- and offsets were then determined as the nearest reversal in the sign of acceleration.

Hand position data were up-sampled to 1 kHz by linear interpolation and filtered using a second-order Butterworth filter (15 Hz cut-off). Hand latency was computed as the first sample exceeding 5% of the peak hand velocity in that trial. Hand movement offset was detected when the finger landed within ±0.80 mm of the screen. If no interception was detected online, interception time and position were determined offline as the maximum hand position in the z-dimension. We inspected all trials manually and excluded trials in which observers blinked or when the eye tracker lost the signal (3.2% of trials across participants).

## Statistical Data Analyses

Our results are based on the analyses of three dependent variables. (1) We calculated estimated target speed from the 2D hand interception position. For each individual trial, we determined which target trajectory best fitted the observed interception position, as follows. We simulated 600 target trajectories with launch speeds ranging from 0.1°/s to 60°/s in 0.1°/s steps. We then determined the trajectory that produced the smallest Euclidian distance to the interception position (**Fig.1A**). The corresponding target speed that best fit to the observed interception position was labelled the estimated target speed for that trial. (2) The same analysis was repeated using the eye position at the time of interception. (3) We analysed the amplitudes of the detected saccades during each trial to obtain a readout of audio-visual signal integration at different time points (early catch-up saccade versus later predictive saccades; **Fig. 4A**). Because trajectories showed their strongest differences in their vertical component, we restricted this analysis on the vertical saccade amplitudes. For each dependent variable we applied a within-subject z-score outlier detection (data points were excluded if they were >3 SD from an observer’s mean).

To assess effects of stimulus volume and presentation time on our dependent variables, we calculated observers’ means per condition and fed the data into repeated measures (rm) ANOVA in R [41] with an alpha level of .05. To correct for multiple comparisons within multiway ANOVA, we applied a sequential Bonferroni-correction [42]. Bonferroni-correction is applied to all post-hoc pairwise comparisons (two-sided, paired *t*-tests).

---

**Figure S1.**
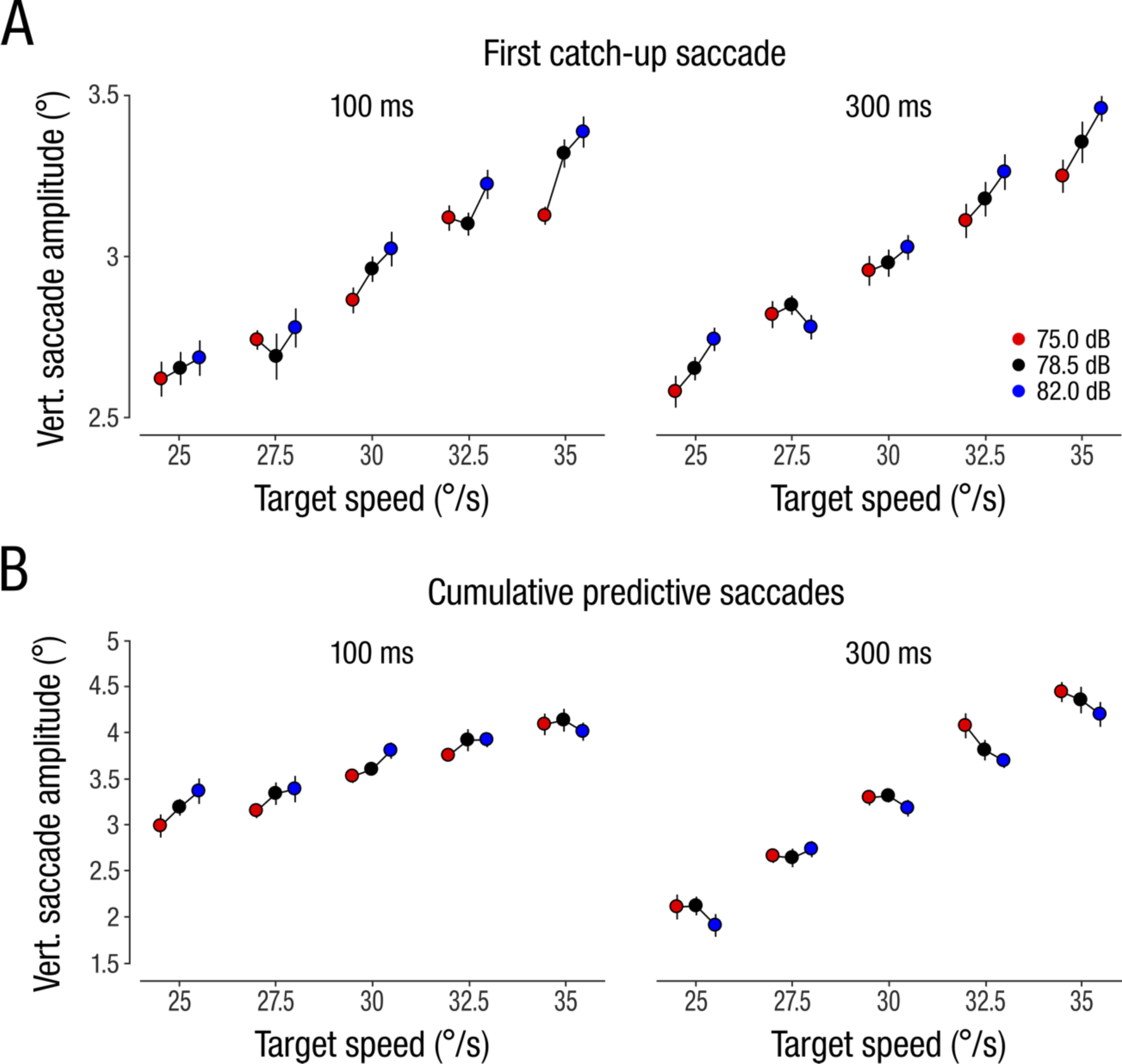
Saccade analyses per target speed condition. **(A)** Effect of sound volume on vertical saccade amplitudes for the first catch-up saccades after target onset, separately for each target speed condition for the 100 ms (left panel) and 300 ms (right panel) presentation times. **(B)** Effect of sound volume on the cumulative vertical amplitudes of all subsequent predictive saccades. Circles and error bars show means ±1 within-subject SEM.

**Figure S2.**
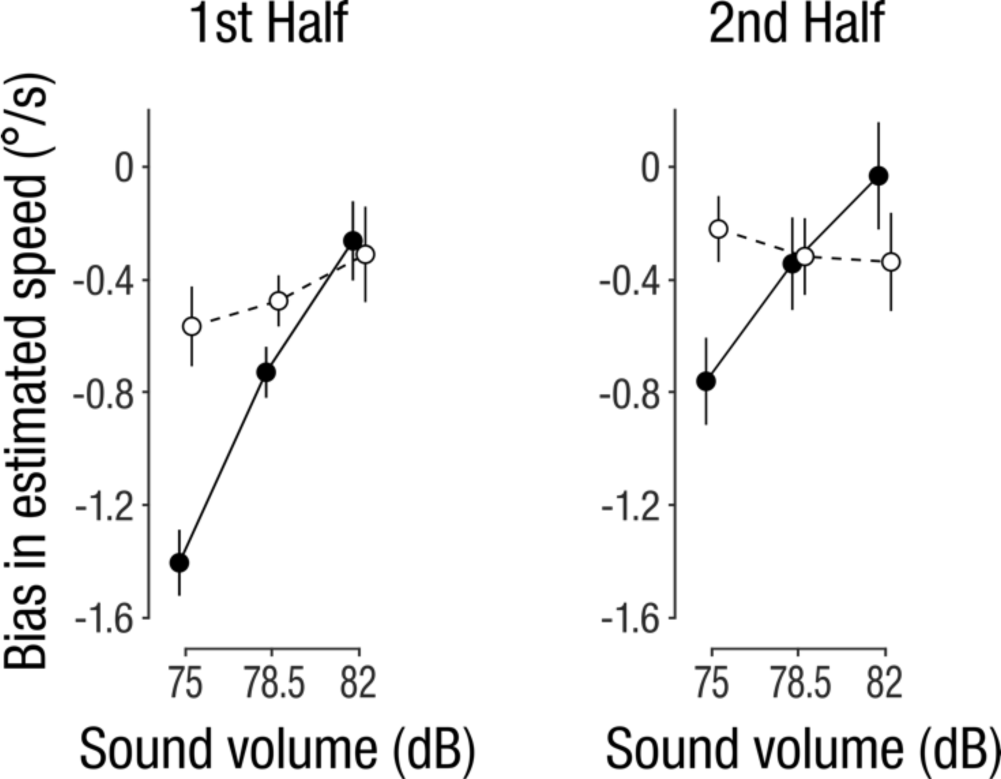
Bias in estimated target speed split separately for the first and second half of the experiment. Solid lines and filled circles: 100 ms, dashed lines and open circles: 300 ms condition. Circles and error bars show mean ±1 within-subject SEM.

